# Bioreactor cultures of brain endothelial cells produce extracellular vesicles with comparable characteristics as that of batch cultures

**DOI:** 10.64898/2026.10.05.756840

**Authors:** Balaji Govindaswamy, Chia-Chen Lu, Nikhila Achanta, Edward Y. Chen, Kristen Rebeles, Paromita Paul Pinky, Vivek Basudkar, Fauzia Parween, Devika S Manickam

**Affiliations:** Department of Neurology, McGovern Medical School, University of Texas Health Science Center at Houston, TX 77030; College of Natural Sciences and Mathematics, University of Houston, Texas, 77004; Department of Pharmaceutical Sciences, Duquesne University, Pittsburgh, PA 15282

**Keywords:** Extracellular vesicles, brain endothelial cells, mitochondria, ATP, bioreactor, batch cultures, large EVs

## Abstract

Our prior works have demonstrated the therapeutic potential extracellular vesicles (**EVs**) derived from batch cultures of brain endothelial cells (**BECs**) owing to their ability to transfer mitochondria to injured recipient cells. However, conventional batch/flask cultures limit scalable and reproducible production of EVs. Here, we evaluated a CELLine AD1000 bioreactor as a platform for sustained production of EVs from mouse b.End3 BECs and compared bioreactor-derived (**BR**) EVs with EVs obtained conventional batch cultures (**BT**). Large EVs (**lEVs**) and small (**sEVs**) were isolated using a sequential centrifugation protocol and were characterized using dynamic light scattering, nanoparticle tracking analysis, microBCA assay, western blotting, proteomics analysis, and their ability to rescue bioenergetics in oxygen-glucose deprived (**OGD**) BECs. Bioreactor cultures maintained high cell viabilities (∼90%) for up to eight weeks and produced approximately three- to four-fold greater EV protein yields than batch cultures. BR EVs showed reproducible particle characteristics consistent with EV subtypes derived from BT cultures. CD63, CD9 and Arf6 were detected in EV fractions, whereas the mitochondrial outer-membrane protein TOMM20 was enriched in lEVs. Proteomic analyses revealed that bioreactor-derived lEVs retained the mitochondrial cargo signature observed in batch-derived lEVs. Both bioreactor and batch EVs restored relative ATP levels in ODG-exposed BECs. These findings suggest that bioreactor BEC cultures are a scalable approach for generating functionally active BEC-EVs for downstream characterization and therapeutic development.

**Graphical abstract:** 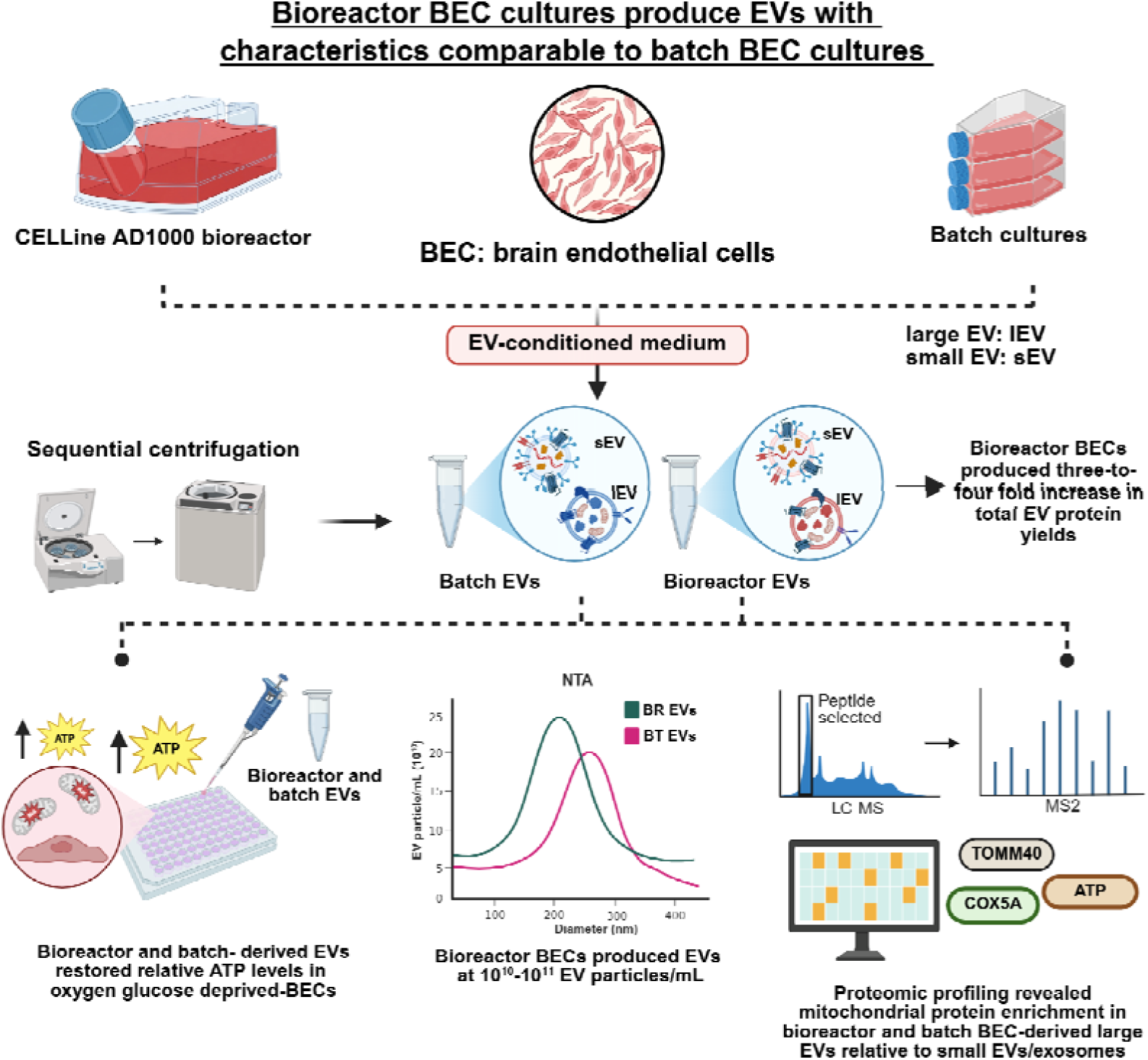

## Introduction

Extracellular vesicles (**EVs**) are a heterogenous population of lipid bilayer-enclosed particles released by all cell types and are widely recognized as integral components of intercellular communication (Zhu *et al*. 2026). They are broadly categorized into microvesicles/large EVs that are shed from the plasma membrane and exosomes, that are derived via endosomal biogenesis pathways although operational definitions remain heavily dependent on isolation method and the analytical framework used (Budgude *et al*. 2025). EVs are biologically important because they transport bioactive protein, lipids, metabolites, nucleic acids, and mitochondria (Nguyen *et al*. 2022; Mo *et al*. 2025; Vanherle *et al*. 2025), allowing them to influence recipient cell phenotype through paracrine, endocrine, and local microenvironmental signaling pathways (Abuna *et al*. 2026; Saviana *et al*. 2026).

EVs can modulate key biological processes including inflammation, angiogenesis, immune regulation, extracellular matrix remodeling and cellular metabolic adaptation (Xiang *et al*. 2025). Owing to these effects, EVs are increasingly being explored as promising nanocarriers for regenerative medicine, immunomodulation, and targeted drug delivery (Li *et al*. 2022). Our laboratory has studied EVs originating from brain microvascular endothelial cells (**BECs**) as a novel carrier for mitochondria (D’Souza *et al*. 2021; Dave *et al*. 2024). We have previously demonstrated that large EVs (>200 nm in particle diameter) can transfer mitochondria cargo into recipient oxygen-glucose deprived (**OGD**) BECs, where the mitochondria colocalize with the recipient mitochondrial network and increase relative ATP levels and mitochondrial function. Interestingly, the mitochondria-specific effects were observed in the large- but not small EV/exosome-treated recipient BECs (Dave *et al*. 2022, Dave *et al*. 2024; D’Souza *et al*. 2021; Dave *et al*. 2023). Intravenously injected large EVs resulted in a reduced infarct size and improved sensorimotor functions in a mouse model of transient ischemic stroke, suggesting that delivery of EV mitochondria is a promising approach to protect the post-ischemic BBB (Dave *et al*. 2024). We also demonstrated that formulations of EVs with an exogenous HSP27 protein rescued BEC tight junction integrity after OGD (Dave *et al*. 2023).

In all of our prior works, we isolated BEC-EVs using standard batch cultures in adherent 182 cm^2^ tissue culture flasks (**T175**) because it allowed facile culturing of BECs. However, flask-based production is often constrained by limited surface areas available for large scale BEC culturing, need for growth medium replenishment every other day post-passaging/culture inoculation, and relatively modest EV yields (Park *et al*. 2025). These limitations can be especially significant for BECs, which require stable culture conditions to maintain phenotype and vesicle output over time. The WHEATON CELLine AD1000 bioreactor supports long-term culture in a two-chamber format and has been shown to improve EV production efficiency relative to conventional flask systems (Artuyants *et al*. 2021; Hisey *et al*. 2022). The upper compartment maintains nutrient exchange through a semipermeable membrane, while the lower compartment allows accumulation of EV-rich conditioned medium with minimal dilution. This configuration not only reduced the frequency of medium replacement/replenishment but also created a physiologically relevant microenvironment with enhanced cell-to-cell interactions and prolonged variability. The EV composition is influenced by cell state, culture density, nutrient gradients, extracellular stress, and duration of conditioning, all of which may differ between a bioreactor and a standard flask (Jeske *et al*. 2022; Shekari *et al*. 2023; Paganini *et al*. 2023). Mitchell *et al*. reported that an Integra CELLine culture system can greatly increase exosome yield from tumor cell cultures while preserving exosome morphology, phenotype, and immune function. The authors compared CELLine cultures with conventional flask cultures and found a 12-fold higher exosome production from mesothelioma cells and about 8-fold higher output from NKL suspension cells, alongside lower maintenance costs (Mitchell *et al*. 2008). Similarly, Hisey *et al*. reported that CELLine AD1000 bioreactors produce large, consistent amounts of EVs from multiple breast cancer cell lines, resulting in an overall resource efficient EV production process than conventional cultures. They also demonstrated that core-EV associated proteins stayed relatively stable over the bioreactor lifetime, monitored by tracking medium volume, shed-cell viability by Trypan blue exclusion assay, and EV particle numbers using NTA; and changes in these parameters were associated with declining culture stability (Hisey *et al*. 2022).

In this study, we cultured mouse b.End3 BECs in an AD CELLine1000 bioreactor as well as in conventional tissue culture flasks to enable a direct comparison of EVs produced from bioreactor *vs*. batch cultures. Bioreactor cultures of BECs were monitored for tracking cell growth kinetics at regular intervals to determine cell expansion and identify optimal culture conditions for EV harvesting. Large as well as small EVs (<200 nm particle diameters) were isolated from both bioreactor and batch cultures using a sequential centrifugation protocol. We compared the characteristics of bioreactor *vs*. batch BEC culture-derived EVs using a combination of assays. Total EV protein content was quantified using a microBCA protein assay, particle diameter, zeta potential and EV concentration were determined using dynamic light scattering and nanoparticle tracking analyses to determine vesicle characteristics. We performed western blotting to identify canonical EV associated markers and completed proteomic profiling to distinguish the cargo composition of bioreactor *vs*. batch culture-derived EVs. Finally, the bioactivity of EVs were determined by measuring changes in relative ATP levels in recipient OGD BECs treated with EVs using a luminescent ATP assay.

## Materials and methods

### Materials

Pierce Micro BCA and BCA protein assay kits were purchased from Thermo Fisher Scientific (Rockford, IL). Cell Titer Glo 2.0 reagent was purchased from Promega (Madison, WI). Dulbecco’s Modified Eagle’s medium (DMEM) and 1X PBS were obtained from Cytiva (Marlborough, MA). Penicillin-Streptomycin was purchased from Invitrogen (Carlsbad, CA). Western electrophoresis buffers were obtained from Research Prospect International (Mount Prospect, IL). WHEATON CELLine AD1000 bioreactors were purchased from DWK life sciences (Mill Ville, NJ). All primary and secondary antibodies were obtained from Abcam (Waltham, MA). Mouse brain endothelial cells (b.End3) were purchased from ATCC (Manassas, VA).

### Cell culture

Mouse brain endothelial (**b.End3**) cells were maintained in T75 tissue culture flasks in a humidified incubator at 37°C with 5% CO_2_. Cells were cultured in complete growth medium consisting of high glucose DMEM supplemented with 10% FBS and 1% Penicillin-Streptomycin. Culture medium was replaced every other day until cells reached 95-100% confluence. Cells between passages 26 and 30 were used in experiments.

### Bioreactor seeding and culture maintenance

To equilibrate the semipermeable membrane and hydrate polyethylene terephthalate layer in the bioreactor, 50 mL of prewarmed growth medium was added to the upper medium compartment of CELLine AD 1000 bioreactor for five minutes (Artuyants *et al*. 2021). b.End3 cells at passage# 27 were seeded into the lower cell compartment at a density of 1.5 x 10^6^ cells/mL in 15 mL growth medium. The upper medium compartment was filled with 500 mL of the same growth medium. Bioreactor was incubated at 5% CO_2_ incubator at 37°C. After an initial adaptation period, culture medium in both the cell and medium compartment was replaced twice weekly. Cell density was monitored by collecting 20 µL medium from cell compartment, mixing it 1:1 with Trypan blue, and quantifying viable cells using a Countess automated cell counter (Invitrogen). Post 30 days of culture, 15 mL of EV-conditioned medium from cell compartment was collected for EV isolation and immediately replaced with an equal volume of fresh growth medium.

### Conventional batch cultures

For comparison with bioreactor cultures, b.End3 cells between passage# 27 to 29 were cultured in 182 cm^2^ tissue culture flask until they reached 90-95% confluency. Post confluency, cells were then washed with 1X PBS and incubated with conditioning medium: DMEM high glucose + 10% exosome-depleted FBS for 48 hours in a humidified 5% CO_2_ incubator at 37°C.

### EV isolation from bioreactor and conventional cultures

EVs were isolated using our previously reported differential centrifugation protocol (Dave *et al*. 2026; Pinky *et al*. 2026) from EV-conditioned medium collected from CELLine AD 1000 bioreactor (**BR**) or conventional batch cultures (**BT**). At approximately 30 days post-seeding, when bioreactor cultures reached 80-90% viability, 15 mL and 60 mL of EV-conditioned medium (20 mL from each T-175 batch culture; three T-175 flasks were used per isolation) were collected from the bioreactor cell compartment and batch cultures, respectively. EV-conditioned medium was collected in polypropylene tubes and centrifuged at 300*x*g for 11 mins at 4°C to remove dead cells. Subsequently, the collected supernatants were further centrifuged at 2000*x*g for 22 mins at 4°C to pellet apoptotic bodies and cell debris using an Allegra X-30R Centrifuge (Beckman Coulter, IN). Following this, the supernatant was transferred into polycarbonate tubes and centrifuged at 20,000*x*g for 45 mins at 4°C to pellet large EVs (**lEVs**) using an Sorvall WX+ UltraSeries Centrifuge (Thermo Scientific, MA). Next, the supernatant was collected and filtered through 0.22 µm sterile PES membrane syringe filter and centrifuged at 120,000*x*g for 70 mins at 4°C to pellet small EVs (**sEVs**) Sorvall WX+ UltraSeries Centrifuge. The isolated EVs were resuspended in sterile 1X PBS and stored at -20°C until further use.

### MicroBCA assay

Total EV protein content was measured using a Pierce MicroBCA assay. Frozen EV samples were thawed on ice and lysed using 1X RIPA buffer. Next, 150 µL of BCA standards and lysed EV samples were mixed with 150 µL of the Micro BCA working reagent (reagent A: reagent B: reagent C at 25:24:1 volume ratio). The plate was then incubated at 37°C for 2 hours. Post incubation, the absorbance was measured at 562 nm using an EnSpire 2300 Multimode Plate Reader (Perkin Elmer, MA).

### Dynamic light scattering

EV particle diameter, polydispersity index and zeta potential were measured using dynamic light scattering (**DLS**) on a Malvern Zetasizer Pro (Malvern Panalytical Inc. Westborough, PA). Samples were prepared at 0.1 mg protein/mL concentration in 1X PBS and 10 mM HEPES buffer at pH 7.4 for particle size and zeta potential measurements, respectively. Average particle diameter, polydispersity index and zeta potential values were reported as mean ± standard deviation (n=3).

### Nanoparticle tracking analysis

EVs particle concentration (particles/mL) was quantified by nanoparticle tracking analysis (**NTA**) using a ZetaView Evolution Twin system (Particle Metrix, Germany). EV samples were diluted 1:2000 in sterile 1X PBS before analysis to obtain an optimal measurement range of approximately 100-200 particles/frame, with particle drift maintained between 0 to 50 µm/s. Measurements were acquired using the 488-nm laser channel with autofocus, camera gain: 55 and shutter speed: 150. Particle traces were analyzed using ZetaSphere software. Data were reported as mean ± standard deviation from three independent measurements (n=3).

### Western blotting

EV-associated protein markers in EVs isolated from bioreactor and batch cultures were studied using western blotting. Briefly, EV lysates comprising 100 µg total protein were mixed with 4x Laemmli buffer. This mixture was heated at 95°C for five minutes using a heating block (Thermo Scientific, MA). The samples and the premixed molecular weight markers (dual color ladder, 250 kDa-10 kDa) were electrophoresed on a 4-15% Mini-PROTEAN TGX Stain-Free Precast Gels (Bio-Rad Laboratories, Inc. CA) at 100-150V for 50 minutes. The proteins were transferred onto a 0.45 µm PVDF membrane using a transfer assembly (Bio-Rad Laboratories, Inc. CA) at 80V and under 300 mA for 70 mins. Next, the membranes were washed with Tris Buffered Saline with 0.1% Tween 20 (**T-TBS**) and were blocked using Bio-Rad EveryBlot blocking buffer solution for 15 minutes at room temperature. The membranes were incubated overnight with either rabbit anti-CD63 (1:1000), rabbit anti-CD9 (1:1000), rabbit anti-Arf6 (1:500) or rabbit anti-TOMM20 (1:500), primary antibody prepared in T-TBS at 4°C. The membrane was washed with T-TBS and incubated with anti-rabbit HRP antibody (0.05 µg/mL) in T-TBS at room temperature for an hour. The blots were washed with T-TBS and then incubated for 5 minutes in a 1:1 solution of SuperSignal West Pico Plus Luminol Enhancer and SuperSignal West Pico Plus Stable Peroxidase (ThermoFisher, MA). The membranes were scanned using a Bio-Rad ChemiDoc imager on the chemiluminescence and colorimetric settings (Bio-Rad Laboratories *Inc*. CA).

### Effects of bioreactor and batch culture derived EVs on relative ATP levels in b.End3 BECs under normoxic and oxygen-glucose deprived (OGD) conditions

To stimulate ischemia *in vitro*, b.End3 cells were exposed to OGD conditions, and the resulting cell viability and relative ATP levels were evaluated by using a Cell Titer Glo ATP assay. b.End3 cells were seeded in 96-well plates at a density of 16,500 cells/well. Confluent b.End3 were incubated for four hours with OGD medium defined as follows: 120 mM NaCl, 5.4 mM KCl, 1.8 mM CaCl_2_, 0.8 mM MgCl_2_, 25 mM Tris-HCl, pH 7.4 for 4 hours in an OGD chamber (Billups Rothenberg, Del Mar, CA) pre-flushed with 5% carbon dioxide, 5% hydrogen and 90% nitrogen at 37°C. For normoxic conditions, the culture plates were incubated in a humidified incubator with 95% air and 5% CO_2_. For normoxic and hypoxic conditions, the medium was removed and the recipient b.End3 cells were incubated with bioreactor or batch culture-derived lEVs and sEVs at doses of 10, 25 and 50 µg EV protein/well in a humidified incubator for 24 hours. EVs were diluted in complete culture medium and OGD medium for normoxic and hypoxic conditions, respectively. Post treatment, cells were replaced with prewarmed growth medium and an equal amount of Cell Titer Glo 2.0 reagent was added. The plates were incubated for 15 mins at room temperature on an orbital shaker in the dark and relative luminescence units were measured at 1s integration time using EnSpire 2300 Multimode Plate Reader (Perkin Elmer, MA). Relative ATP levels were calculated by normalizing the relative luminescence unit (RLU) of the treatment groups to the RLU of control, untreated cells (equation (1)) (Dave, Stolz, and Manickam 2023; Pinky *et al*. 2026; Dave *et al*. 2024, 2026).

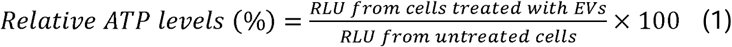

### Proteomic characterization of EVs from bioreactor and batch cultures

EV samples were denatured in 5% SDS, reduced with 5 mM DTT (90°C, 5 min), and alkylated with 10 mM iodoacetamide (15 min, room temperature). Proteins were processed on S-Trap Micro Spin Columns (ProtiFi) per manufacturer protocol and digested overnight at 37°C with a trypsin/LysC mixture. Peptide concentration was determined by colorimetric assay (Pierce Quantitative Colorimetric Peptide Assay, Thermo Scientific) and samples were reconstituted at 100 ng/µL in 0.1% formic acid containing 0.015% DDM. LC-MS/MS analysis was performed on a nanoElute2 UHPLC system coupled online to a timsTOF Ultra 2 mass spectrometer (Bruker Daltonics) using a 25 cm × 75 µm × 1.5 µm PepSep column with a 28-min active gradient. Data were acquired in diaPASEF mode (m/z 400–1000; ion mobility 0.64–1.37 1/K0; 24 DIA windows; cycle time 0.96 s). Raw data were searched in library-free mode using DIA-NN (v2.2.0) against a mouse proteome FASTA (UniProtKB; 17,228 entries; accessed November 2024). Search parameters included trypsin/P digestion (up to 1 missed cleavage), variable oxidation (M) and N-terminal acetylation, fixed carbamidomethylation (C), and precursor FDR set at 1% with match-between-runs enabled. Protein quantification was performed using the iBAQ approach via the gpGrouper algorithm (Saltzman *et al*. 2018), followed by median normalization and log2 transformation. Differential expression was assessed using moderated t-tests implemented in the R package limma with Benjamini-Hochberg FDR correction. Gene set enrichment analysis (GSEA) was performed against MSigDB gene set collections using signed log p-values (differential analysis) or z-scored expression values (ssGSEA) (Liberzon *et al*. 2015; Subramanian *et al*. 2005; Mootha *et al*. 2003).

For unsupervised clustering, log-transformed abundances were z-scored per protein, and samples and proteins were clustered hierarchically by Ward’s method (ward.D2) using Euclidean distance. Mitochondrial proteins were identified using the MitoCarta annotation provided with the dataset, and their over-representation among differentially abundant proteins was tested with a one-sided Fisher’s exact test against all quantified proteins as background (Rath *et al*. 2021). Agreement between culture formats was assessed by Pearson correlation of lEV vs. sEV log2 fold changes in BR and BT across proteins detected in all samples. Agreement between culture formats was assessed at the proportion of proteins significant in either within-format lEV vs. sEV comparison whose log2 fold change had the same sign in BR and BT, and by Pearson correlation of lEV *vs*. sEV log2 fold changes in BR and BT across proteins detected in all samples.

### Statistical analysis

Data are presented as mean ± SD from at least three independent measurements. Statistical analyses were performed using GraphPad Prism software (version 10; San Diego, CA). Comparisons between two groups were performed using an unpaired two-tailed student’s t-test. Comparisons among multiple groups were analyzed using one-way or two-way analysis of variance (ANOVA), as appropriate, followed by Dunnett’s multiple comparison test. A p-value of < 0.05 was considered statistically significant.

## Results

### EVs isolated from bioreactor cultures retain physiochemical characteristics of batch culture-derived EVs

Efficient and sustained cell expansion is an important prerequisite for obtaining EV-enriched medium over extended periods. b.End3 cell growth and viability were monitored longitudinally after seeding in the CELLine AD1000 bioreactor. As shown in (**Fig. 1A**), b.End3 cells maintained high viability during the early and mid-phases of culture, with approximately 80-90% live cells observed for 1-7 weeks and we start to notice a decline in % viability post 7 week. In parallel, cell density progressively increased from approximately 10^4^-10^5^ cells/mL from the initial seeding to approximately 2x10^6^ cells/mL at later time points, suggesting substantial expansion of BECs within the bioreactor. Cell density remained elevated through the mid-culture phase 3-4 week, whereas a modest decline in viability was observed toward the end of the culture window around 8^th^ week. To further examine the morphological status of cells in the bioreactor, 15 mL of culture medium from cell compartment was collected and transferred to T75 culture flask. The recovered cells displayed BEC morphology shown in (**Fig. S1A**). Culture medium from bioreactor and conditioned medium from batch cultures was processed through differential centrifugation to isolate lEVs and sEVs (**Fig. 1B**). Total protein yield for BR and BT EV subsets was quantified using a microBCA assay (**Fig. 1C**). BR lEVs yielded 0.3-0.4 mg protein/mL of EV-conditioned medium across six independent isolations. In comparison, BT lEVs showed approximately 0.1 mg protein per mL of EV-conditioned medium, demonstrating a three-to-four-fold increase in BR EV yields. BR sEVs yielded 0.4-0.5 mg/mL, whereas BT sEVs ranged 0.07-0.1 mg/mL, the BR sEVs approximately showed a 4-fold greater yield compared to batch cultures. These findings indicate that the CELLine AD 1000 bioreactor enabled a greater yield and therefore, increased recovery of EVs.

**Figure 1:** EV isolation from AD CELLine 1000 bioreactor and batch cultures **A**. Growth kinetics of AD CELLine 1000 cultures over two months post-seeding. Weekly EV isolations started 30 days post-seeding. **B**. Schematic of differential ultracentrifugation workflow for isolating lEVs and sEVs from conditioned medium. **C**. total protein yield per mL of EV-conditioned medium from six bioreactor (BR1-BR6) and three batch cultures (BT1-BT3) EV preparations determined using a microBCA assay. **D**. Particle diameters of EVs isolated from bioreactor and batch cultures across independent isolations. **E**. Surface charge/ζ-potential of EVs isolated from bioreactor and batch cultures across independent isolations. **F**. NTA analysis of EVs. **G**. Western blotting analysis of TOMM20, CD63, ARF6 and CD9 expression in BR and BT EVs. Data represented as mean ± SD; statistical significance was determined by one way-ANOVA with multiple comparisons (*p<0.05, **p<0.01, ***p<0.001, ****p<0.0001; ns, not significant).

We next compared the physicochemical characteristics of EVs derived from bioreactor *vs*. batch cultures. BR lEVs ranged 150-190 nm in Z-average diameter, whereas BR sEVs were consistently smaller at 40-70 nm (**Fig. 1D**). BT lEVs were both larger with diameters ranging from 230-320 nm, while BT sEVs showed particle diameters ranging from 40-55 nm. These values were similar to BR sEVs, suggesting that the BEC culturing format alters the lEV particle diameters to a marked extent compared to sEVs. BT lEVs were approximately 52% larger than BR lEVs based on the particle diameters, while BT sEVs were approximately 12% smaller than BR sEVs. As expected, zeta potentials stayed negative across all EV groups regardless of culturing conditions, with BR sEV and BT lEV ranging from nearly -10 to -30 mV (**Fig. 1E**).

EV particle concentrations of BR and BT EVs were determined using NTA (**Fig. 1F**). lEV concentrations ranged from 1.8×10¹¹ ± 6.1×10□ particles/mL in BR isolations 1 and 2, with BR isolations 3–6 showing concentrations of 8.3×10¹□, 1.2×10¹¹, 1.2× 10¹¹, and 9.3×10¹□ particles/mL, respectively. sEV concentrations were both lower and noticeably tighter across the series, ranging from 6.52×10¹□ ± 3.4×10□ particles/mL (isolation 1) to 1.0×10¹¹ ± 3.9×10□ particles/mL (isolation 5), with isolations 2, 3, 4, and 6 falling in between these two values (7.0×10¹□, 8.2×10¹□, 8.4×10¹□, and 9.6×10¹□ particles/mL). Similarly, BT lEVs concentrations increased across the passage series (27-29), from 1.3x10¹□ ± 1.1x10^8^ particles/mL at P27, 1.5x10^10^± 9.3x10^7^ particles/mL at P28 and 2.3x10¹□ ± 1.1x10^8^ particles/mL at P29. In contrast, BT sEV concentrations were consistently higher than lEVs ranging from 5.3x10^10^ to 4.2x10^10^ particles/mL. These findings are consistent with the microBCA data, which showed a greater EV protein recovery from the bioreactor cultures. Overall, these findings indicate that bioreactor cultures generated consistently greater concentrations of both EV subtypes, with yields generally maintained within 10^10^-10^11^ particles/mL range across six consecutive isolations.

We performed western blotting analysis for BR and BT EV subtypes using the following markers: TOMM20, translocase of the outer mitochondrial membrane, tetraspanins CD63 and CD9, and Arf6 that regulates the release of plasma membrane-derived microvesicles (Muralidharan-Chari *et al*. 2009). TOMM20 (16 kDa) was detected in both lEVs (BR and BT) but was absent in sEVs. This finding suggests that mitochondria or mitochondrial proteins are selectively incorporated into the lEV fraction and is excluded from the sEVs. In contrast, CD63 resolved a broad band spanning approximately (30-60 kDa), consistent with its characteristic heterogenous glycosylation and was detected at comparable intensities across BR and BT EVs, supporting earlier findings that CD63 is expressed in both EV subsets. CD9 (25 kDa) was similarly detected across all lanes, though signal intensity was consistently higher in BR EVs relative to BT EVs (Jhaveri *et al*. 2024). ARF6 were detected in BR and BT EV subtypes supporting the presence of plasma membrane-derived lEVs albeit low levels of expression were also seen in sEVs (**Fig. 1G**).

### Serum EVs contribute to 20% of total bioreactor EV yields

FBS used in bioreactor cultures to maintain BECs is a source of bovine EVs. To apply a background correction, we subtracted serum EV protein yields from BR EV protein yields (**Fig. 2E**). We isolated serum EVs from 20 mL complete culture medium (DMEM high glucose supplemented with 10% FBS) through sequential centrifugation (**Fig. 2A**). We observed lEV and sEV protein yields of 0.08-0.1 mg/mL (**Fig. 2B**). Serum lEVs measured 38-58 nm, barely larger than standard sEV diameters, and serum sEVs clustered tightly at 31-33 nm, the observed smaller size distribution of serum EVs likely reflects a mixture of lipoproteins and protein aggregates (**Fig. 2C**), with zeta potentials ranging -9 to -16 mV (**Fig. 2D**). For background correction, we used serum lEV and sEV concentrations of 0.09 and 0.08 mg/mL, respectively. The corresponding BR yields were 0.32 mg/mL for lEV and 0.4 mg/mL for sEV. Subtracting the serum EVs background from BR EVs resulted in adjusted/true BR yields of 0.24 mg/mL for lEVs and 0.31 mg/mL for sEVs, respectively. To summarize, the serum EVs that are considered as background contributed to 24% of total bioreactor EVs.

**Figure 2:** **A**. Schematic representation of serum EV isolation from growth medium. **B**. Total protein yield of lEVs and sEVs isolated from four tubes, demonstrating comparable recovery across isolations. **C**. DLS analysis of particle diameters of serum-derived EVs **D**. Surface charge characterization of serum-derived EVs **E**. Serum EV background correction from bioreactor EVs. Data are presented as mean ± SD; statistical significance was determined by one-way ANOVA with multiple comparisons (*p<0.05, **p<0.01, ***p<0.0001; ns, not significant).

### Bioreactor and batch derived EVs restore relative ATP levels in oxygen-glucose deprived (OGD) BECs

Functional activity of BR and BT derived EVs were determined by measuring relative ATP levels in recipient OGD BECs using CellTiter-Glo 2.0 assay, which is a bioluminescence-based readout of relative ATP levels. Normoxic BECs were first treated at 10, 25 and 50 µg EV protein/well. Most treatment groups showed no significant deviation in relative ATP levels to untreated cells; both BR lEVs and BT sEVs at the highest dose tested (50 µg protein/well) produced a modest but statistically significant reduction, while remaining within approximately 85-95% of the untreated cells (**Fig. 3A**). OGD was then applied as an established *in vitro* model of cerebral ischemic injury, in which combined oxygen and glucose deprivation results in loss of mitochondrial function and consequently, ATP depletion (D’Souza *et al*. 2021a; Dave *et al*. 2024; Dave *et al*. 2023). As reported in our previous studies, OGD-treated recipient BECs showed 40% of the ATP levels, confirming a robust bioenergetic deficit suitable for assessing EV mediated rescue. Treatment with lEVs or sEVs from either BR or BT cultures produced a statistically significant restoration of ATP at every dose tested (p<0.0001 relative to untreated OGD cells), with recovered ATP levels ranging from 65% to 85% of the normoxic baseline (**Fig. 3B**). To determine whether this rescue effect was retained by BR EVs, each of the six individual BR lEVs or sEVs were evaluated at the medium dose of 25 µg protein/well, alongside BT lEVs and sEVs (**Fig. 3C**). All treatment groups produced statistically significant ATP recovery relative to untreated OGD cells.

**Figure 3:**
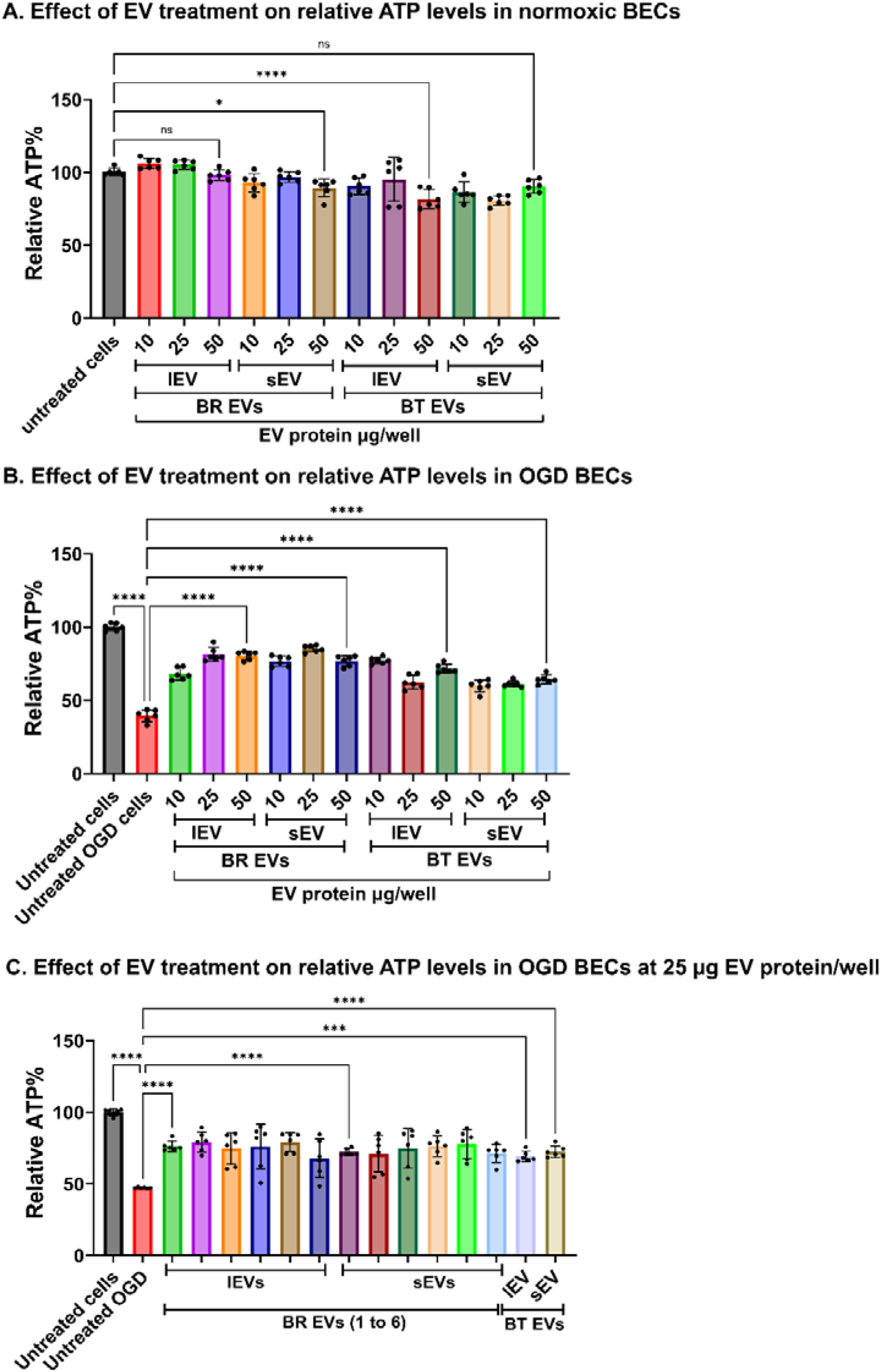
**A**. Relative cellular ATP levels in normoxic BECs treated with bioreactor-derived EVs (BR EVs) or batch culture-derived EVs (BT EVs) at 10, 25, and 50 µg EV protein per well compared with untreated cells. **B.** Relative ATP levels in hypoxic BECs following treatment with BR-derived or BT-derived EVs at increasing EV protein concentrations of 10, 25, and 50 µg/well. **C.** Comparison of ATP levels in hypoxic BECs treated with 25 µg/well of BR and BT EVs with untreated OGD controls. Data are presented as mean ± SD; statistical significance is indicated by the corresponding significance bars and was determined by one-way ANOVA (*p<0.05, ****p<0.0001; ns, not significant).

### Proteomic profiles distinguish bioreactor- and batch-derived EVs

Quantitative LC-MS/MS proteomics on lEV and sEV fractions from three BR isolations and three BT preparations were performed to compare the cargo of BR- and BT-derived EVs, quantifying 4,712 proteins. Unsupervised hierarchical clustering of all quantified proteins separated samples first by culture format (BR or BT format), with all BR samples clustering apart from all BT samples (**Figure 4**). Within each culture format, lEV and sEV fractions from the same preparation clustered together in some cases (BR1 and BT P29), indicating that preparation-specific effects may have contributed to sample similarity alongside EV subset. Generally, BR samples lended higher protein identification than in BT samples (mean ∼3,300 vs. ∼2,300 protein groups per sample; **Supplementary Table 1**), and many proteins detected in BR samples were not detected in BT samples, visible as non-detected regions in **Figure 4**.

**Figure 4:**
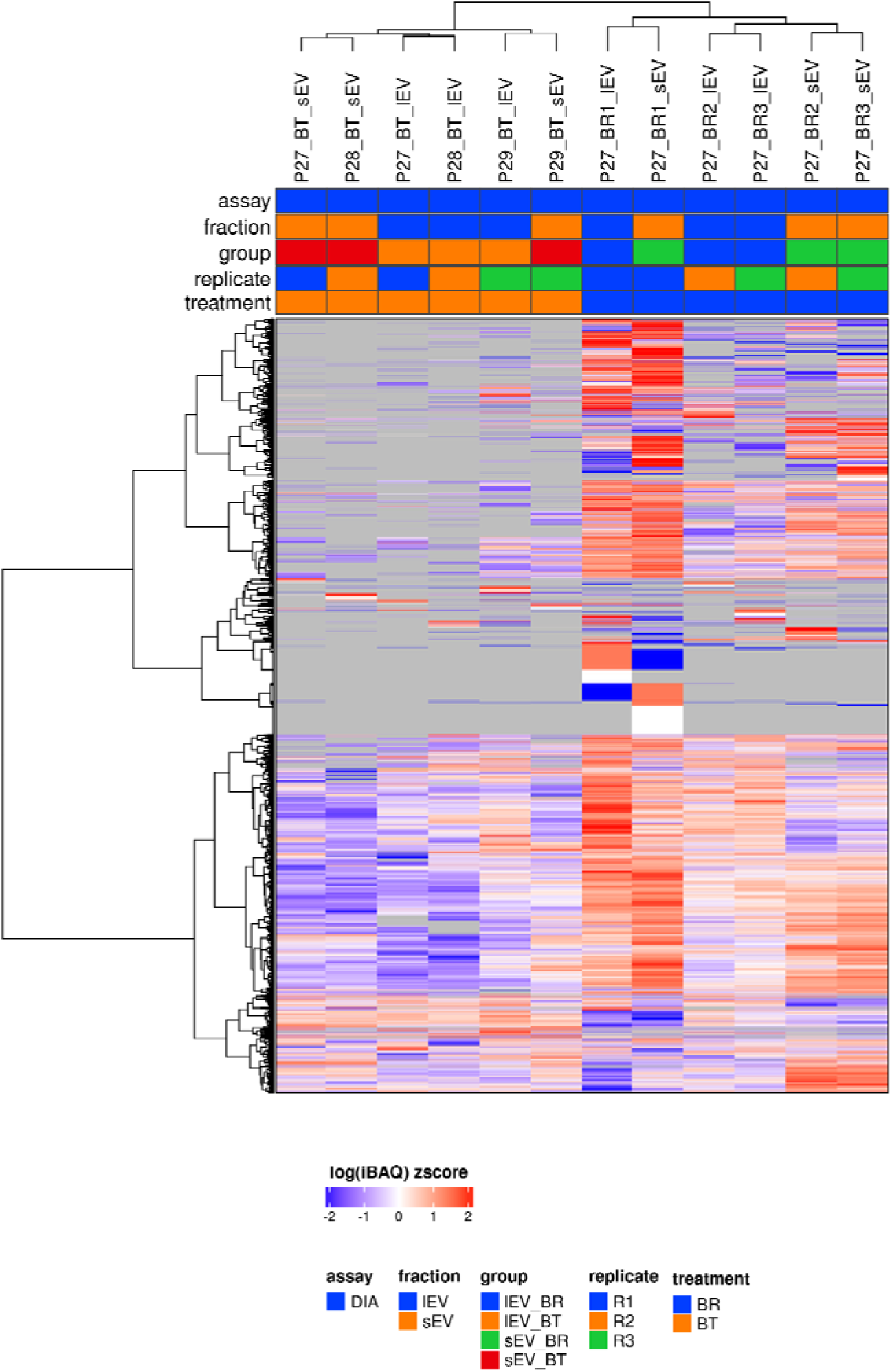
Global proteomic profiles of bioreactor- and batch-derived EVs. Unsupervised hierarchical clustering of all quantified proteins (n = 4,712) in lEV and sEV fractions from BR and BT cultures. Values are median-normalized, log-transformed iBAQ intensities, z-scored per protein. Red indicates higher and blue lower relative abundance; gray indicates proteins not detected in that sample. Rows and columns were clustered by Ward’s method (ward.D2). BR-EVs: three consecutive EV isolations from a single bioreactor culture; BT-EVs: independent EV preparations from passages 27–29.

### Mitochondrial proteins are enriched in lEVs from both bioreactor and batch cultures

Consistent with the clustering in **Figure 4**, differential abundance analysis identified 280 proteins differing between BR and BT lEVs and 960 between BR and BT sEVs, of which, a large majority were more abundant in BR EVs (**Figure 5A, 5B**). Approximately half of these differences (138/280 proteins and 468/960 proteins) are detected in at least two of three replicates in both culture formats; the remaining proteins fell below the detection threshold in BT samples, so these differences may partly reflect the greater detection depth of the BR samples. In contrast, the differences between EV fractions were conserved across culture formats. lEVs and sEVs differed by 38 proteins in BR and 24 in BT EVs (**Figure 5C, D**).

**Figure 5:**
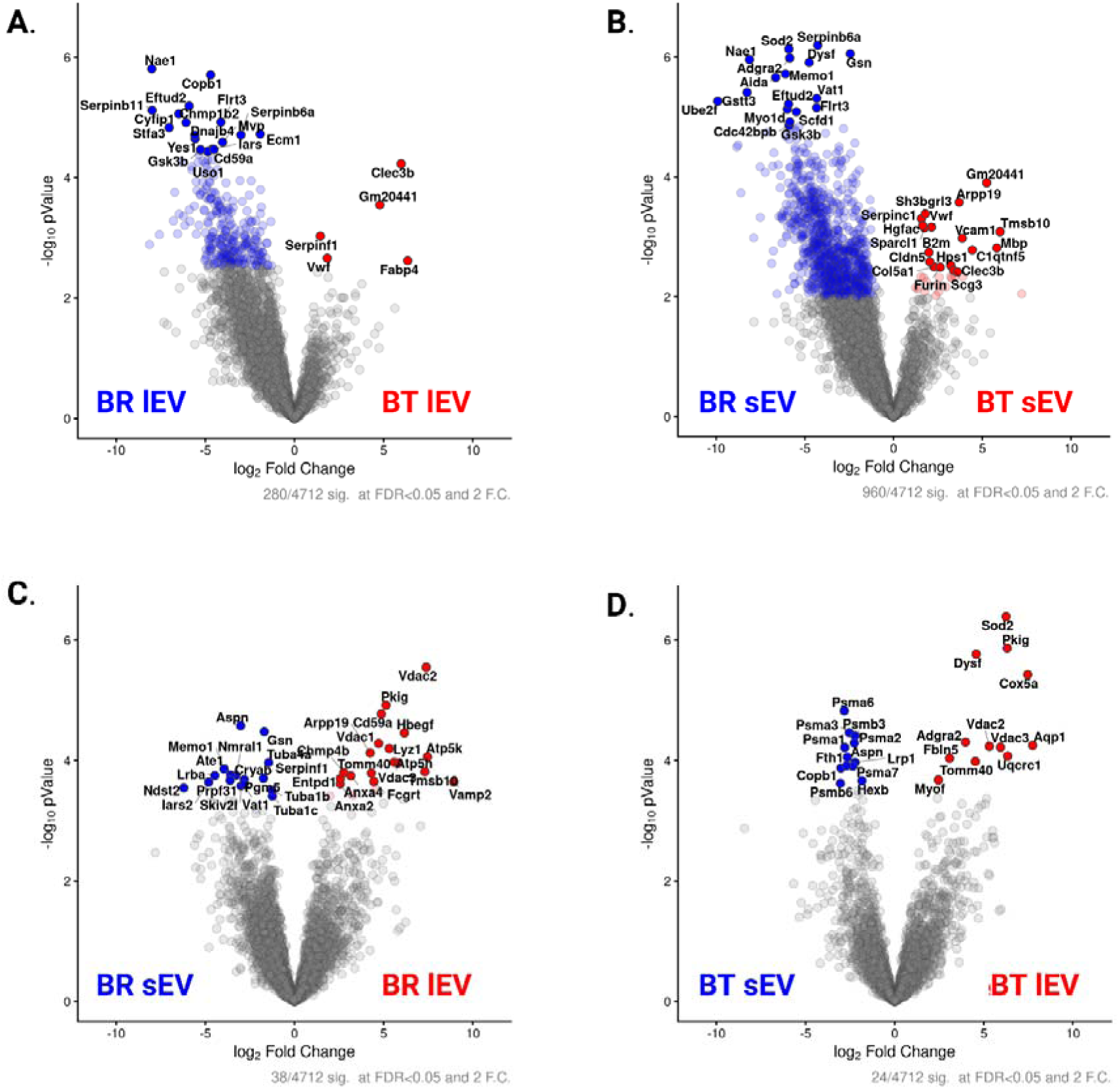
Differential protein abundance in bioreactor- and batch-derived EVs. Volcano plots from quantitative LC-MS/MS (diaPASEF) proteomics comparing **A.** BR *vs*. BT lEVs, **B.** BR *vs*. BT sEVs, **C.** lEVs *vs*. sEVs within BR cultures, and **D.** lEVs *vs*. sEVs within BT cultures. Each point represents one protein (4,712 quantified). The x-axis shows log₂ fold change; the y-axis shows −log₁₀ p-value from limma moderated t-tests fitted to a four-group model. Colored points meet Benjamini–Hochberg FDR < 0.05 and ≥2-fold change: blue, enriched in BR EVs (A, B) or sEVs (C, D); red, enriched in BT EVs (A, B) or lEVs (C, D). Gray points are not significant; selected top-ranked proteins are labeled. Significant proteins: (A) 280, (B) 960, (C) 38, (D) 24. BR-EVs are three consecutive EV isolations from a single bioreactor culture; BT-EVs are three independent preparations (passages 27–29).

Additionally, lEV-enriched proteins often included mitochondrial proteins across both culture formats, such as VDAC isoforms, TOMM40, and ATP synthase subunits in BR, and VDAC2/3, TOMM40, COX5A, UQCRC1, and SOD2 in BT. Many were significantly overrepresented among lEV-enriched proteins in both BR (5 of 21, 24%; p=0.01) and BT (6 of 12, 50%; p<0.001) relative to their frequency among all quantified proteins (6.6%), but not among sEV-enriched proteins. The lEV *vs*. sEV differences were congruent between culture formats, with 81% (46 of 57) of proteins significant in either comparison changed in the same direction in BR and BT, and fold changes were correlated across proteins detected in all samples (r=0.49). Together with the mitochondrial TOMM20 enrichment in lEVs confirmed using western blotting (**Figure 1G**), these data indicate that bioreactor culture preserves the mitochondrial cargo signature of lEVs, despite global proteomic differences between culture formats.

## Discussion

In the present study, we established and evaluated a bioreactor-based workflow for continuous isolation of EV subtypes (lEVs *vs*. sEVs) from BECs over eight weeks. We observed a consistent EV recovery across independent bioreactor isolations, with reproducible protein yields, particle concentration, particle size distributions, zeta potentials, EV-associated protein markers, and functional activity in an OGD model.

Bioreactors offer an efficient approach for EV scale up by supporting high cell density and repeated EV harvest. The CELLine AD1000 is a two-compartment bioreactor that supports high density adherent cell culture through continuous nutrient exchange and waste removal. By retaining secreted EVs in a small cell compartment, it enables repeated production of BEC-EVs suitable for downstream functional studies. We maintained the CELLine AD 1000 bioreactor cultured with b.End3 BECs at high density and high viability ranging from 80-95% for up to eight weeks until the bioreactor showed a decline in total cell viability and live cells (**Fig. 1A**). The bioreactor design features a small, high-density cell compartment layered with a polyethylene terephthalate matrix for adherent cells to attach and expand, covered with 10 kDa cellulose acetate-semipermeable membrane which allows continuous flow of nutrients and metabolites and allows continuous culturing without the need to passage cells. Hisey *et al*. showed that the use of CELLine AD1000 supported long-term culture of multiple breast cancer cell lines and enabled consistent EV collection across bioreactor culture, although EV yield and cargo profiles varied among cell lines and culture period. (Hisey *et al*. 2022). Similarly, Fu *et al*. demonstrated that serum-free CELLine AD1000 culture supports reproducible EV production from ovarian cancer cell lines for downstream characterization and biomarker studies (Fu *et al*. 2026). Our findings extend this evidence to b.End3 BECs by showing sustained cell viability, reproducible EV yield and physicochemical characteristics, and maintained functional activity in a OGD model. <u>This is</u> <u>an important advance that demonstrates that hard-to-culture BECs can be cultured in a</u> <u>bioreactor format allowing continuous isolation of EVs</u>.

We observed a reproducible total protein yield recovery from BR lEVs and sEVs across six independent/consecutive isolations (**Fig. 1C**). The average BR protein concentrations ranged from 0.28-0.37 mg/mL for lEVs and 0.34-0.45 mg/mL for sEVs from 15 mL of EV-conditioned medium in comparison to BT EVs where the average lEV protein yield was 0.07-0.08 mg/mL and sEV showed 0.07-0.1 mg/mL across independent isolations using 60 mL of conditioned medium. These results indicate that the bioreactor can generate sufficient EVs for characterization and repeated functional studies, with greater yields than conventional batch cultures. The greater recovery observed in our study is consistent with the rationale that high-density culture platforms and a smaller cell compartment increase EV output by accommodating a larger number of viable cells in a confined culture volume. Fuzeta *et al*. evaluated a microcarrier-based Vertical-Wheel bioreactor for production of human mesenchymal stromal cell-derived EVs and reported a 5.7-fold increase in EV concentration in conditioned medium and an approximately three-fold increase in EV production per cell compared with conventional static T-flask cultures. These findings align well with our observations that bioreactor cultures can increase EV recovery relative to standard culture systems (De Almeida Fuzeta *et al*. 2020). Similarly, Costa *et al*. studied the production of stromal cell EVs in stirred-tank bioreactors. They found that growing cells in a bioreactor raised the yield of EVs by about 2.2 times compared with cells grown on tissue culture surfaces/2D cultures. They also reported that fine-tuning the cell-culture environment and collecting the EVs after collection had an effect on production rates on EVs (Costa *et al*. 2023). Although our system, the type of cells, the protocol we use for collecting the EVs and the measure we use are different from those used by Costa *et al*., both studies show that using a bioreactor can boost EV production when compared with batch cultures. Gobin *et al*. further reported that human bone marrow mesenchymal stromal cells cultured in a hollow-fiber bioreactor produced higher EV yields that retained an immunomodulatory signature related to the donor cells (Gobin *et al*. 2021).

We observed substantially higher protein recovery from the bioreactor cell compartment, which contained growth medium with cultured cells. To determine the extent to which bovine EVs contributed background to BEC-EVs, we isolated serum EVs. The mean protein yield was 0.05-0.08 mg/mL for serum lEV and 0.07-0.09 mg/mL for serum sEV per 20 mL of medium (**Fig. 2B**). We then normalized the bioreactor yields (BR4 to BR6) to 20 mL, the normalized yields were subtracted with serum EVs, the adjusted BR yields were 0.24 and 0.31 mg/mL for BR lEV and sEV, indicating that 76% of the bioreactor EVs are BEC-EVs, whereas the rest are bovine EVs (**Fig. 2E**). Noticeably, Lehrich *et al*. reported that FBS-derived EVs persist even after depleting exosomes from FBS, indicating that a serum-EV background is unavoidable (Lehrich *et al*. 2018). Consistent with this finding, Pham *et al*. detected approximately 5.3 x 10^7^ bovine EVs/mL and showed that residual bovine EVs were present in human cell derived EV fractions, even when the serum was used at only 0.5% concentration (Pham *et al*. 2021).

We next ran DLS analysis for EV fractions from both from BT and BR cultures. DLS measures the hydrodynamic diameter of particles in suspension by analysing fluctuations in scattered light. Consistent with our prior findings (Dave *et al*. 2022, Dave *et al*. 2024; D’Souza *et al*. 2021; Dave *et al*. 2023), we observed that the lEV fraction had larger Z-average diameters compared to sEV fractions across both BR and BT conditions (**Fig. 1D**). In the BR EVs, lEVs generally exhibited particle diameter of approximately 155-190 nm, whereas sEVs were smaller with a Z-average diameters of approximately 40-65 nm. Consistent with findings of Eitan *et al*. who reported that most membrane vesicles isolated from serum measured around 30-120 nm (Eitan *et al*. 2015), the serum EVs (**Fig. 2C**) displayed a smaller diameter range of 30-50 nm. The BT lEVs showed a heterogenous distribution ranging from 250 to 300 nm. The obtained particle diameters are consistent with standard particle diameters of lEVs ranging more than 200 nm and below 1000 nm, whereas our sEVs ranged from 30-150 nm, consistent with MISEV guidelines (Welsh *et al*. 2024). We performed NTA to quantify EV particle/mL obtained from BR and BT cultures. Our data showed that BR lEVs ranged from 7x10^10^ ± 8x10^8^ to 1.8x10^11^ ± 6.1x10^9^ particles/mL and the BR sEV particle concentration ranged between 6.5x10^10^ ± 3.4x10^9^ to 1.0x10^11^ ± 3.9x10^9^, whereas BT lEVs ranged from 1.7x10^10^ ± 5.2x10^9^ and sEVs ranged 4.5x10^10^ ± 6.6x10^9^ (**Fig. 1F**). Overall, NTA results demonstrated that the BR workflow enabled repeated recovery of concentrated EV fractions with particle concentrations consistently maintained within the 10^10^-10^11^ particles/mL range across independent isolations.

Our western blotting analysis showed the presence of the EV-associated tetraspanins CD63 and CD9 in the BR and BT cultured derived EV fractions (**Fig. 1G**). The detection of these tetraspanins supports the interpretation that the isolated fractions contained EVs. We also observed differences in CD63 and CD9 band intensity between lEV and sEV fractions and between BR and BT derived fractions. This variability is expected because EVs are heterogenous, and the relative abundance of individual tetraspanins can vary according to the cell state and EV biogenesis pathway. Kowal *et al*. demonstrated CD9, CD63 and CD81 for EV subpopulations and reported that tetraspanins should not be interpreted as universal markers for single homogenous EV subtype (Kowal *et al*. 2016). ARF6 marker was strongly detected in BR EVs, and a weak signal was seen in BT EVs, supporting the presence of plasma-membrane associated vesicles in both the EV fractions. We also detected TOMM20, an outer mitochondrial membrane protein with a more prominent signal in the lEV fraction (**Fig. 1G**). This observation is important because lEVs represent the larger particle fraction that packs intact mitochondria in vesicles during biogenesis (Dave *et al*. 2026; Dave *et al*. 2023; Dave *et al*. 2024).

We evaluated the functional activity of BR and BT EVs, our ATP results demonstrate that BR derived EVs were well tolerated under normoxic conditions and provided partial protection against ATP depletion during hypoxia/OGD stress (**Fig. 3A**). Under hypoxic conditions, untreated cells showed substantial decline in ATP levels, whereas treatment with both BR lEVs and sEVs increased ATP levels across the tested dose range (**Fig. 3B**). The consistent response observed at 25 µg protein/well across six independent BR isolations suggests that the bioreactor workflow generated EVs with reproducible bioactivity. Thus, BR and BT EV-treated cells exhibited similar relative ATP levels under OGD conditions (**Fig. 3C**). Our results suggest that bioreactor culturing format sustains healthy cells that release EVs with ATP-increasing activity while enabling repeated EV collection at higher rates of recovery. D’Souza *et al*. demonstrated that BEC-derived EVs increased ATP levels in recipient BECs subjected to OGD in a dose-dependent manner and identified 14.6 µg protein/well as a minimum effective dose for ATP rescue post-OGD (D’Souza *et al*. 2021). Likewise, Dave *et al*. reported that mitochondria containing BEC-derived large EVs increased ATP levels and improved mitochondrial function in OGD-exposed BECs (Dave *et al*. 2023).

We performed quantitative proteomics on lEV and sEV fractions from BR and BT cultures to determine the effects of culture format on the packaging of EV cargo. Global protein profiles differed between culture formats: samples clustered primarily by culture format (**Fig. 4**), and several hundred proteins differed in abundance between BR and BT EVs, most of them higher in BR EVs (**Fig. 5A, B**). Part of this difference reflects detection depth, as BR samples yielded approximately 1.5-fold more protein identifications than BT samples (**Table S1**), and about half of the differentially abundant proteins were not detected in BT samples. Differences in EV composition between culture formats are nonetheless expected, as culture conditions such as cell density, nutrient exchange, and conditioning time influence EV cargo, and Hisey *et al*. reported that cargo profiles of bioreactor-derived EVs varied with cell line and culture period (Hisey *et al*. 2022).

## Limitations

Bioreactor b.End3 BEC cultures showed a decline in cell viability 45 post-days of culture suggesting that bioreactor cultures have a defined lifetime that allowed six subsequent EV isolations. BECs were maintained in complete culture medium during the lifetime of the bioreactor and therefore, the isolated BEC-EVs were not excluded from serum EVs that were accounted as background EVs in our study. The luminescence-based ATP assay does not allow measuring absolute concentrations of cellular ATP levels in EV-treated cells but rather the luminescent readouts from the assay have been calculated and expressed as %relative ATP levels. We recognize that changes in relative ATP levels is only one way to measure the bioenergetic response in EV-treated cells. Our future studies will determine the effects of EV treatments on rescuing mitochondrial function in recipient OGD BECs using a Seahorse extracellular flux assay. We are currently optimizing the experimental setup to determine basal mitochondrial function in OGD-exposed b.End3 BECs prior to measuring the effects of EV treatment.

## Conclusions

Findings from this study suggest that CELLine AD 1000 bioreactor is a scalable approach for producing BEC-EVs. Bioreactor cultures increased total EV protein yields relative to conventional flask cultures while preserving key vesicle characteristics and ATP-restorative abilities in oxygen-glucose deprived recipient BECs. Proteomic profiling further showed that, although global protein profiles differed between culture formats, bioreactor-derived lEVs retained the mitochondrial cargo signature observed in batch-derived lEVs. These findings support bioreactor cultures as a promising approach for generating BEC-EVs for further characterization and functional studies.

## Author contributions

**Balaji Govindaswamy**: Writing – review and editing, Writing – original draft, Visualization, Methodology, Investigation, Formal analysis, Data curation. **Chia-Chen Lu**: Investigation, Methodology, Formal analysis, Data curation. **Nikhila Achanta**: proteomic data analysis, methodology, writing – review and editing. **Edward Y. Chen**: Investigation, Data curation. **Kristen Rebeles**: Methodology, Formal analysis, Data curation, Investigation. **Paromita Paul Pinky**: Methodology, Investigation. **Vivek Basudkar**: Investigation. **Fauzia Parween**: Methodology. **Devika S Manickam**: Writing – review and editing, Writing – original draft, Validation, Supervision, Resources, Project administration, Methodology, Funding acquisition, Formal analysis, Data curation, Conceptualization.

## Acknowledgements

The authors gratefully acknowledge National Institutes of Health grant number 7R01NS136752-02 for funding support. Devika S Manickam is grateful for a UT System Translational STARs Award that partly supported the procurement of the NTA system used in this study. We thank Dr. Colin Hisey (Northwestern University) for his protocol guidance in establishing the bioreactor cell cultures. We acknowledge the services from Baylor College of Medicine’s Mass Spectrometry Proteomics Core led by Dr. Anna Malovannaya and Mr. Antrix Jain along with the support of Drs. Kwangwon Lee, Mei Leng, and Alexander Saltzman and Ms. Shirley Wang. The core is supported by CPRIT Core Facility Award (RP210227), P30 Cancer Center Support Grant (NCICA125123), Intellectual Developmental Disabilities Research Center Award (P50 HD103555), Gulf Coast Center for Precision Environmental Health (P30 ES030285), and S10 High End Instrument Award (OD026804).

## List of frequently used non-standard abbreviations

BCA: bicinchoninic acid
BECs: brain endothelial cells
BR EVs: bioreactor EVs
BT EVs: batch EVs
EVs: extracellular vesicles
FBS: fetal bovine serum
lEVs: large EVs
RLU: relative luminescence unit
sEVs: small EVs

