## Supplemental File for "Bioreactor cultures of brain endothelial cells produce extracellular vesicles with comparable characteristics as that of batch cultures"

6431 Fannin Street, MSE R274

Houston, TX 77030

ORCID: 0000-0002-6363-3679

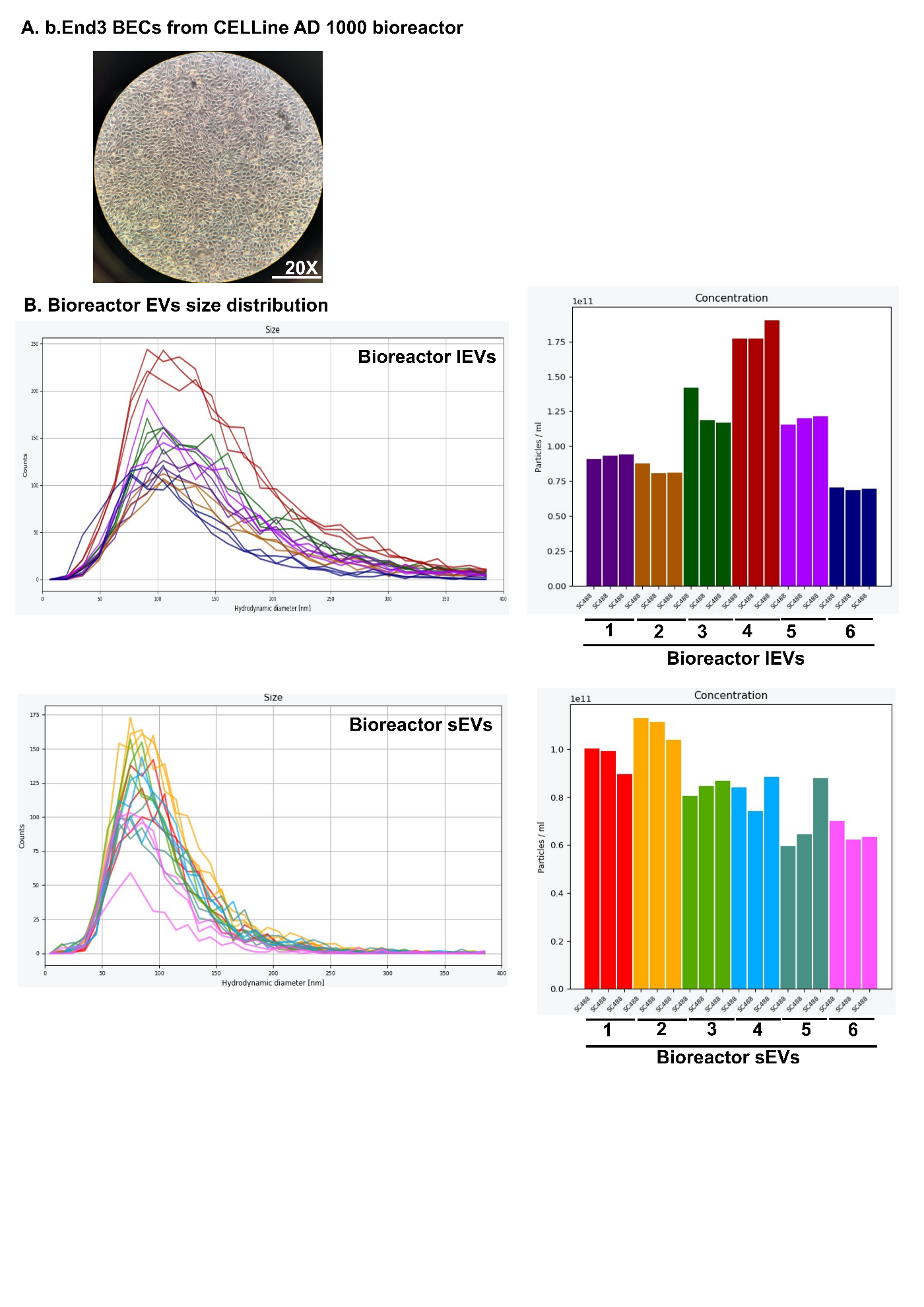

**Supplementary figure 1:** **A.** Representative brightfield image of cells harvested from CELLine AD1000, scale bar 20x magnification. **B.** Nanoparticle tracking analysis of BR lEV and sEVs with representative particle diameter distribution and particle concentration measured for n=3 replicates.

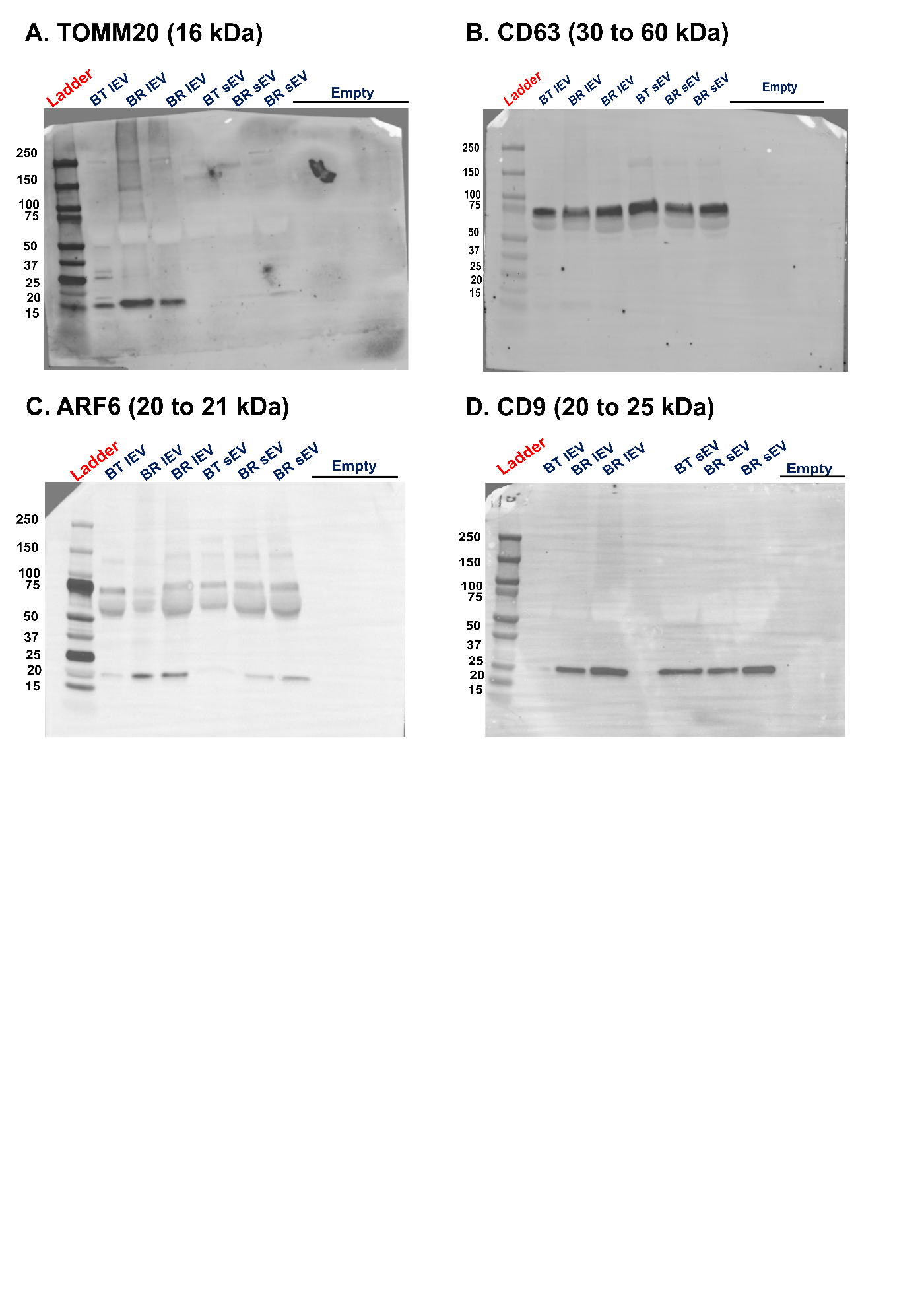

**Supplementary figure 2:** Uncropped images of western blots. **A.** EV fractions stained for TOMM20 (1:1000), **B.** CD63 (1:1000). **C.** ARF6 (1:500) **D.** CD9 (1:2000).

**Supplementary Table 1** reports identification depth for each proteomics sample. Bioreactor EV (**BR**) samples yielded more identifications than batch EV (**BT**) samples (mean 3,432 *vs*. 2,298 protein groups; 1.49-fold), while lEV and sEV fractions from the same preparation were measured at similar depth. These data support interpretation of the culture-format comparisons in **Figures 4 and 5**, where differences involving proteins not detected in BT samples partly reflect detection depth.

| **Sample** | **Culture format** | **EV fraction** | **Preparation** | **Protein groups** | **Peptides** | **PSMs** |
| --- | --- | --- | --- | --- | --- | --- |
| P27_BR1_lEV | BR | lEV | BR1 | 3,909 | 25,395 | 27,620 |
| P27_BR1_sEV | BR | sEV | BR1 | 4,066 | 26,727 | 29,096 |
| P27_BR2_lEV | BR | lEV | BR2 | 2,963 | 15,962 | 17,334 |
| P27_BR2_sEV | BR | sEV | BR2 | 3,182 | 17,584 | 19,098 |
| P27_BR3_lEV | BR | lEV | BR3 | 3,156 | 17,507 | 18,995 |
| P27_BR3_sEV | BR | sEV | BR3 | 3,318 | 19,394 | 21,108 |
| P27_BT_lEV | BT | lEV | BT P27 | 2,082 | 9,797 | 10,527 |
| P27_BT_sEV | BT | sEV | BT P27 | 2,212 | 10,963 | 11,897 |
| P28_BT_lEV | BT | lEV | BT P28 | 2,201 | 10,319 | 11,098 |
| P28_BT_sEV | BT | sEV | BT P28 | 2,078 | 9,935 | 10,796 |
| P29_BT_lEV | BT | lEV | BT P29 | 2,715 | 13,003 | 14,142 |
| P29_BT_sEV | BT | sEV | BT P29 | 2,497 | 12,687 | 13,754 |

**Supplementary Table 1**: Number of protein groups, peptides, and peptide-spectrum matches (PSMs) identified in each lEV and sEV sample from BR and BT cultures by LC-MS/MS (diaPASEF; DIA-NN, 1% FDR). Protein groups were assigned by gpGrouper. BR samples are three consecutive EV isolations (BR1–BR3) from a single bioreactor culture at passage 27; BT samples are independent batch-culture preparations from passages 27–29. lEV and sEV fractions from the same preparation were isolated from the same conditioned medium. Summary rows give mean values per culture format and the BR/BT ratio.

**Supplementary Table 2: Proteins differing between lEVs and sEVs in bioreactor and batch cultures.** Table 2 lists the proteins distinguishing lEVs from sEVs in each culture format. Of 57 proteins significant in either format, 46 (81%) changed in the same direction in BR and BT, and five (VDAC2, VDAC3, TOMM40, PKIG, and ASPN) were significant in both. Mitochondrial proteins, including VDAC isoforms, TOMM40, and ATP synthase and respiratory chain subunits, were predominantly lEV-enriched, supporting conservation of the lEV mitochondrial cargo signature across culture formats.

| **Gene symbol** | **BR log2 FC (lEV/sEV)** | **BR adj. p** | **BT log2 FC (lEV/sEV)** | **BT adj. p** | **Significant in** | **Enriched in BR** | **Enriched in BT** | **Same direction** | **MitoCarta** |
| --- | --- | --- | --- | --- | --- | --- | --- | --- | --- |
| Vdac2 | 7.39 | 8.96E-03 | 5.31 | 2.21E-02 | Both | lEV | lEV | Yes | MOM |
| Vdac3 | 5.59 | 3.83E-02 | 5.93 | 2.21E-02 | Both | lEV | lEV | Yes | MOM |
| Pkig | 5.14 | 2.51E-02 | 6.31 | 2.49E-03 | Both | lEV | lEV | Yes |  |
| Tomm40 | 4.31 | 3.83E-02 | 4.52 | 2.81E-02 | Both | lEV | lEV | Yes | MOM |
| Aspn | -3.03 | 2.67E-02 | -2.66 | 2.71E-02 | Both | sEV | sEV | Yes |  |
| Atp5k | 7.46 | 3.83E-02 | 5.17 | 1.30E-01 | BR only | lEV | lEV | Yes |  |
| Aqp1 | 3.43 | 2.31E-01 | 7.74 | 2.21E-02 | BT only | lEV | lEV | Yes |  |
| Atp5h | 5.85 | 3.83E-02 | 4.86 | 7.86E-02 | BR only | lEV | lEV | Yes | MIM |
| Cox5a | 2.41 | 2.51E-01 | 7.46 | 4.27E-03 | BT only | lEV | lEV | Yes | MIM |
| Tmsb10 | 7.31 | 3.83E-02 | 1.39 | 9.32E-01 | BR only | lEV | lEV | Yes |  |
| Uqcrc1 | 2.02 | 3.93E-01 | 6.34 | 2.71E-02 | BT only | lEV | lEV | Yes | MIM |
| Vdac1 | 4.72 | 3.45E-02 | 3.46 | 9.71E-02 | BR only | lEV | lEV | Yes | MOM |
| Lyz1 | 5.32 | 3.68E-02 | 1.77 | 6.31E-01 | BR only | lEV | lEV | Yes |  |
| Sod2 | 0.71 | 6.23E-01 | 6.25 | 1.29E-03 | BT only | lEV | lEV | Yes | Matrix |
| Hbegf | 6.17 | 2.67E-02 | 0.05 | 1.00E+00 | BR only | lEV | lEV | Yes |  |
| Dysf | 1.09 | 3.51E-01 | 4.57 | 2.49E-03 | BT only | lEV | lEV | Yes |  |
| Loxl1 | 4.49 | 4.24E-02 | 1.03 | 9.05E-01 | BR only | lEV | lEV | Yes |  |
| Arpp19 | 4.25 | 3.83E-02 | 1.08 | 8.03E-01 | BR only | lEV | lEV | Yes |  |
| Anxa5 | 3.29 | 4.83E-02 | 1.93 | 3.26E-01 | BR only | lEV | lEV | Yes |  |
| Fcgrt | 4.45 | 3.83E-02 | 0.57 | 9.94E-01 | BR only | lEV | lEV | Yes |  |
| Anxa4 | 3.16 | 3.83E-02 | 1.84 | 2.64E-01 | BR only | lEV | lEV | Yes |  |
| Anxa2 | 2.56 | 3.86E-02 | 2.40 | 7.18E-02 | BR only | lEV | lEV | Yes |  |
| Entpd1 | 2.58 | 3.83E-02 | 1.68 | 1.88E-01 | BR only | lEV | lEV | Yes |  |
| Myof | 1.71 | 1.20E-01 | 2.45 | 4.46E-02 | BT only | lEV | lEV | Yes |  |
| Fbln5 | 0.93 | 4.22E-01 | 3.08 | 2.71E-02 | BT only | lEV | lEV | Yes |  |
| Chmp4b | 2.77 | 3.83E-02 | 0.93 | 6.91E-01 | BR only | lEV | lEV | Yes |  |
| Mfge8 | 1.99 | 4.83E-02 | 0.70 | 7.17E-01 | BR only | lEV | lEV | Yes |  |
| Hexb | -0.08 | 9.39E-01 | -1.84 | 4.46E-02 | BT only | sEV | sEV | Yes |  |
| Tuba1c | -1.25 | 4.83E-02 | -0.87 | 1.90E-01 | BR only | sEV | sEV | Yes |  |
| Tuba1b | -1.27 | 4.24E-02 | -0.90 | 1.72E-01 | BR only | sEV | sEV | Yes |  |
| Tuba4a | -1.45 | 3.83E-02 | -1.10 | 1.09E-01 | BR only | sEV | sEV | Yes |  |
| Nmral1 | -3.28 | 3.83E-02 | -0.11 | 1.00E+00 | BR only | sEV | sEV | Yes |  |
| Skiv2l | -3.00 | 3.89E-02 | -0.44 | 9.86E-01 | BR only | sEV | sEV | Yes |  |
| Psma3 | -0.91 | 2.96E-01 | -2.56 | 2.21E-02 | BT only | sEV | sEV | Yes |  |
| Cryab | -2.82 | 3.83E-02 | -0.77 | 8.23E-01 | BR only | sEV | sEV | Yes |  |
| Psma2 | -1.35 | 1.23E-01 | -2.27 | 2.21E-02 | BT only | sEV | sEV | Yes |  |
| Lrp1 | -1.46 | 1.18E-01 | -2.23 | 2.81E-02 | BT only | sEV | sEV | Yes |  |
| Psma7 | -1.34 | 1.74E-01 | -2.36 | 2.93E-02 | BT only | sEV | sEV | Yes |  |
| Psmb3 | -1.50 | 9.23E-02 | -2.23 | 2.21E-02 | BT only | sEV | sEV | Yes |  |
| Prpf31 | -3.61 | 3.83E-02 | -0.17 | 1.00E+00 | BR only | sEV | sEV | Yes |  |
| Psma6 | -1.14 | 2.19E-01 | -2.84 | 1.40E-02 | BT only | sEV | sEV | Yes |  |
| Fth1 | -1.34 | 2.23E-01 | -2.70 | 2.93E-02 | BT only | sEV | sEV | Yes | unknown |
| Ate1 | -3.59 | 3.83E-02 | -0.61 | 9.65E-01 | BR only | sEV | sEV | Yes |  |
| Psmb6 | -1.43 | 2.68E-01 | -3.05 | 4.72E-02 | BT only | sEV | sEV | Yes |  |
| Psma1 | -1.85 | 1.12E-01 | -2.81 | 2.21E-02 | BT only | sEV | sEV | Yes |  |
| Memo1 | -3.96 | 3.83E-02 | -1.40 | 6.52E-01 | BR only | sEV | sEV | Yes |  |
| Vamp2 | 8.93 | 3.83E-02 | -1.95 | 9.08E-01 | BR only | lEV | sEV | No |  |
| Cd59a | 4.87 | 2.51E-02 | -0.33 | 9.99E-01 | BR only | lEV | sEV | No |  |
| Adgra2 | -1.14 | 4.19E-01 | 3.97 | 2.21E-02 | BT only | sEV | lEV | No |  |
| Gsn | -1.70 | 2.67E-02 | 0.13 | 9.99E-01 | BR only | sEV | lEV | No |  |
| Serpinf1 | -1.75 | 3.83E-02 | 0.15 | 9.99E-01 | BR only | sEV | lEV | No |  |
| Vat1 | -2.88 | 3.86E-02 | 0.61 | 9.26E-01 | BR only | sEV | lEV | No |  |
| Pgm5 | -2.83 | 3.86E-02 | 0.08 | 1.00E+00 | BR only | sEV | lEV | No |  |
| Copb1 | 0.01 | 9.93E-01 | -3.02 | 3.00E-02 | BT only | lEV | sEV | No |  |
| Lrba | -4.47 | 3.83E-02 | 0.72 | 9.70E-01 | BR only | sEV | lEV | No |  |
| Iars2 | -4.82 | 3.83E-02 | 0.42 | 9.99E-01 | BR only | sEV | lEV | No | Matrix |
| Ndst2 | -6.21 | 4.05E-02 | 0.01 | 1.00E+00 | BR only | sEV | lEV | No |  |

**Supplementary Table 2**: All proteins differentially abundant between lEVs and sEVs (Benjamini–Hochberg adjusted p < 0.05 and ≥2-fold change) in either BR or BT cultures (union of **Figures 5C and 5D**; n = 57). Log2 fold changes are lEV relative to sEV (positive, higher in lEVs), from limma moderated t-tests in a four-group model. "Significant in" indicates whether the protein met both thresholds in BR, BT, or both. "Same direction" indicates that the fold change has the same sign in both culture formats, regardless of significance in the other format. MitoCarta indicates mitochondrial sub-compartment annotation (**MOM**, mitochondrial outer membrane; **MIM**, mitochondrial inner membrane; Matrix); blank indicates no mitochondrial annotation.
